# C57BL/6 Mouse Substrain Differences in the Ontogeny of Auditory Fear

**DOI:** 10.64898/2025.12.17.694992

**Authors:** Hanista Premachandran, Kanza Naveed, Maithe Arruda-Carvalho

## Abstract

Early life adversity is a major risk factor for mental disorders later in life. Critically, genetic and individual differences modulate the impact of early life trauma and stress on adult mental health, emphasizing the importance of understanding how the early onset of individual differences may shape behavioral outcomes across life. In rodents, genetic differences can be studied using inbred rodent strains or substrains. While C57BL/6 mouse substrains differ in their adult fear behavior, it is unclear when in development these differences emerge. Here we compared the ontogenesis of remote auditory fear retrieval, spontaneous recovery and renewal in the mouse C57BL/6J (Jax, from Jackson Laboratories) and C57BL/6NTac (Taconic Biosciences) substrains spanning postnatal days (P)16, P18, P21, P25 and P30. Taconic male and female mice show an earlier onset of persistent auditory fear memory and spontaneous recovery (females only) compared to Jax mice. Male Taconic mice also display earlier onset of renewal compared to Jax males. These data identify ontogenetic differences in fear processing among C57BL/6 mouse substrains, highlighting how individual differences can contribute to altered behavioral phenotypes in early development.

## Introduction

The ability to appropriately regulate fear responses is crucial for one’s well-being and survival. Fear regulation differs between individuals^1,2^, with genetic differences noted as a significant source for this variability^3–6^. Individual differences in fear expression are relatively stable over time^7,8^, and are associated with risk for anxiety- and fear-related disorders^9–11^. Importantly, genetic variation also modulates the long-term effects of adverse childhood and adolescent experiences^12–16^, emphasizing the importance of understanding the potential impact of early life individual differences on future mental health outcomes. Although the literature acknowledges individual differences in fear and anxiety-related disorders, rodent research has largely focused on the “average” rodent^2^. This is mostly due to the use of inbred strains believed to be genetically homogenous^17^. Contrary to this perception, decades of research point to substantial differences in fear processing between mouse^9,18–27^ and rat^28–32^ strains, with further evidence of divergent fear and anxiety-related responses in animals from the same mouse strain but acquired through different vendors^19,20,33^, referred to as substrain differences.

The commonly used C57BL/6 mouse strain originated from Jackson Laboratories, but the lack of rederivation within other commercial rodent providers led to genetic drift giving rise to C57BL/6 mouse substrains^19,34,35^. For example, the C57BL/6NTac (Taconic, derived from the 6N mouse line or National Institutes of Health, NIH) and C57BL/6J (Jackson laboratories or 6J) substrains differ by 12-point mutations^36^. This genetic variability is accompanied by differences in contextual fear expression in adulthood^19,26,37–39^, with mice derived from 6N (Taconic) lines showing higher levels of freezing compared to the 6J (Jax) line^19^. While mouse strain and substrain differences in fear processing are well established in adult animals, *when* in development these differences emerge is unknown.

Fear responses change across development in both humans and rodents^40–45^. Specifically, infant rodents display an inability to form life-long or persistent fear memories, also known as infantile amnesia^46–50^. Adult and young rodents also differ in the way they process fear extinction. While extinction in adults typically leads to a return of fear through mere passage of time (spontaneous recovery) or after a change in context (renewal), extinction in younger rodents leads to permanent fear suppression^42–44,51^. Rats^52–55^ and mice^56^ differ in the timing of their transitions toward adult-like fear behavior, but it is unclear whether this extends to ontogenetic variations among rodent strains and/or substrains. Here, we elucidated the ontogenetic profiles for persistent auditory fear memory, spontaneous recovery and renewal among two commonly used mouse substrains: C57BL/6J (Jackson Laboratories) and C57BL/6NTac (Taconic Biosciences). We found that C57BL/6NTac mice display an earlier onset of adult-like fear behavior compared to C57BL/6J mice. These data suggest that, similar to humans, individual differences in mouse fear processing emerge as early as infancy, highlighting the power of using mouse substrains as a tool to probe the neural basis of individual differences in behavior across the lifespan.

## Methods

### Animals

Male and female C57BL/6J (Jackson Laboratories, strain #000664) and C57BL/6NTac (Taconic Biosciences, model #B6-M and B6-F) mice were bred at the University of Toronto Scarborough and kept on a 12h light-dark cycle (lights on at 7:00 A.M.), with access to food and water ad libitum. Mice used for behavioral experiments were of F1 or F2 generation (no noted generation differences between substrains). Animals were housed in plastic Allentown cages (31cm x 17cm x14cm) with paper nesting materials and a plastic igloo, and cages were changed once a week by vivarium staff. For all animals, P0 marked the first day of birth. Mice were weaned at P21 and separated into cages with same-sex siblings (2-4 per cage). All experiments occurred during the light cycle and were performed under the approval of the Animal Care Committee at the University of Toronto.

### Behavior

#### Fear training

Mice were placed in stainless steel chambers (31cm x 24cm x 21cm) from Med Associates (MED-VFC-SCT-M) during training in context A, consisting of a metal grid floor, low intensity house light, and the scent of 70% ethanol. After 2 minutes of habituation to the chamber, mice received 6 pairings of a pure tone (conditioned stimulus, CS: 80dB; 10 seconds in duration) and a footshock (aversive unconditioned stimulus, US: 0.6 mA; 1 second duration) separated by 1-minute intervals^50,56^. The chamber was cleaned with 70% ethanol in between trials. Immediately after training, mice were returned to their home cages.

#### Extinction

One day following fear training, mice were placed in a novel context B, which consisted of round and white plexiglass walls and white floors with no distinct smell. Following a two-minute habituation period, mice were presented with 12 tones (80dB; 10 seconds in duration) separated by 1-minute intervals. Chambers were cleaned with distilled water in between testing.

#### Spontaneous Recovery Testing

One week after extinction training, mice were returned to the extinction context (context B). Following a two-minute habituation period, mice were presented with 5 tones (80dB; 10 seconds in duration) separated by 1-minute intervals. Chambers were cleaned with distilled water in between testing.

#### 9-day Fear Retrieval

To examine the onset of persistent auditory fear memory, mice were placed in context B 9 days after fear training^56^ (no extinction). At the retrieval test, following a two-minute habituation period, all mice were presented with 5 tones (80dB; 10 seconds in duration) separated by 1-minute intervals. Chambers were cleaned with distilled water in between testing.

#### Behavioral analysis

Freezing quantification was automated using Video Freeze software with a motion threshold of 18 and a minimum freeze duration of 1 second as in our previous work^50,57^. For the persistent memory experiments, we analyzed average freezing at CS1-2 of retrieval for direct comparison with Gogolla et al. (2009)^56^ Analysis of average freezing across CS1-5 yielded similar overall differences between age groups for Jax mice (see **Supplementary Table S1** and **Figure S1**), with the exception of P21 Jax males showing less spontaneous recovery than P30 Jax males, and P18 Taconic females showing less spontaneous recovery than P30 females (compared to CS1-2; **Supplementary Table S1**).

### Statistical analysis

Behavioral data are presented as mean ± standard error of the mean (SEM). Graphs and analyses were performed using GraphPad Prism versions 7.05 and 10. Repeated measures (RM) two-way ANOVA was used to assess differences in freezing across tone presentations for fear training and fear retrieval (baseline was excluded from reported analyses). One- and two-way ANOVAs were used to assess the effects of age and/or sex for experiments comparing means of multiple age-groups or rearing conditions. Sexes were collapsed only when there was no significant effect of sex.

Given the absence of sex differences in the persistent auditory fear experiments (**Figure 1**; Jax – Two-way ANOVA: F_interaction 3,45_ = 0.14, *p* = 0.94, F_sex 1,45_ = 0.59, *p* = 0.45, F_age 3,45_ = 15.37, *p* < 0.0001. Taconics – Two-way ANOVA: F_interaction 3,48_ = 0.38, *p* = 0.77, F_sex 1,48_ = 0.92, *p* = 0.45, F_age 3,48_ = 3.51, *p* = 0.022), we collapsed the data across sexes in that dataset. Data were considered statistically significant when the *p* value was less than 0.05. Group differences were assessed through Tukey’s *post hoc* correction test for all comparisons when appropriate.

**Figure 1.**
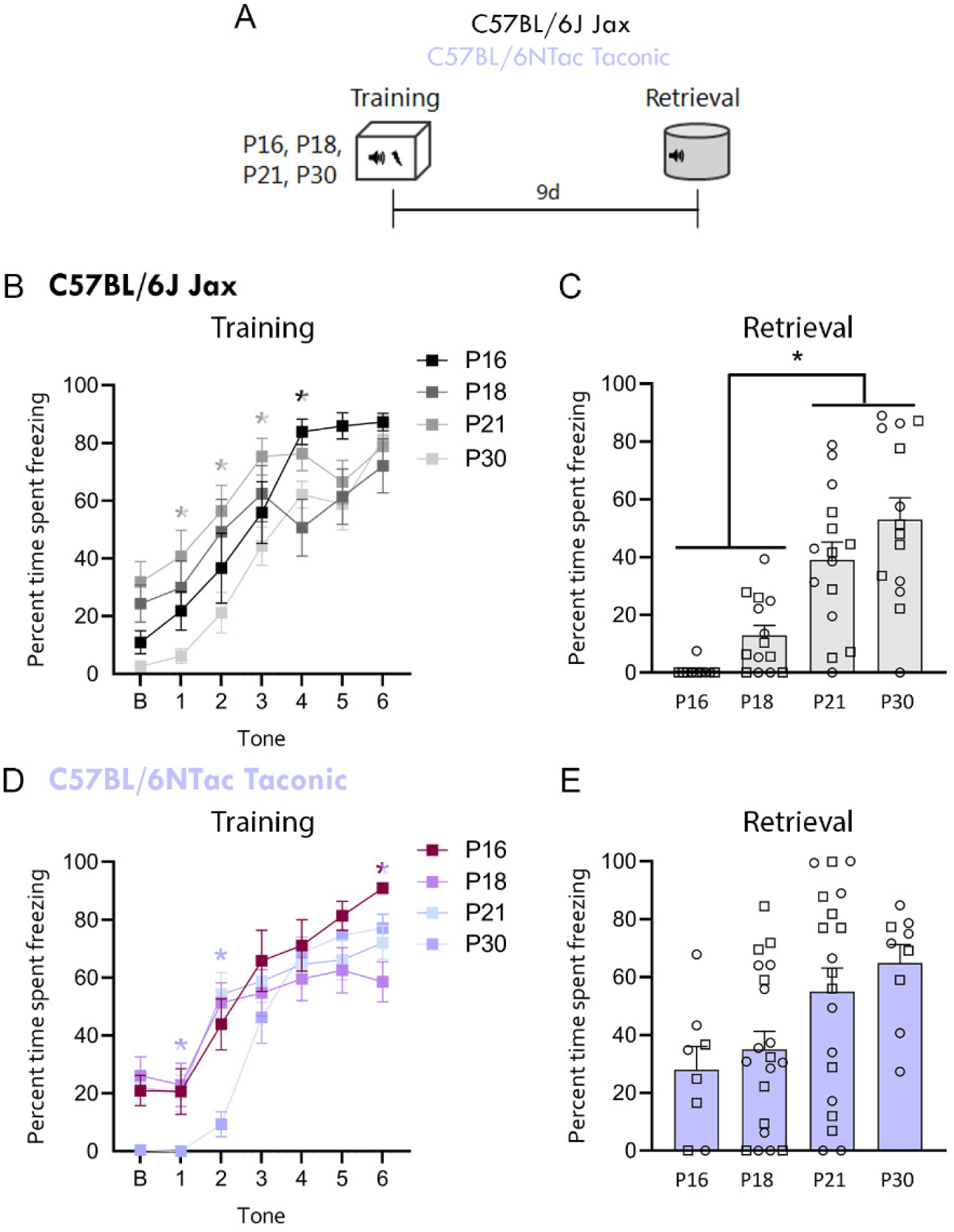
Onset of persistent auditory fear memory in C57BL/6J (Jax) and C57BL/6NTac (Taconic) mice. **A.** Schematic of behavioral design. Mice were fear trained and tested for long-term (9-day) auditory fear retrieval. **B-C**. C57BL/6J (Jax) mice**. B.** Fear acquisition. * represents a difference between P21 vs P30 (tones 1-3), * between P16 vs P18 and P30 (tone 4). **C.** 9-day auditory fear retrieval. **D-E.** C57BL/6NTac (Taconic) mice**. D.** Fear acquisition. * represents a difference between P18, 21 vs 30 (tone 1), * between P30 vs. P16, 18 and P21 (tone 2), * between P16 vs P18, P21 (tone 6). **E.** 9-day auditory fear retrieval. Jax: P16: n = 10, P18: n = 14 , P21: n = 15, P30: n =14. Taconic: P16: n = 8, P18: n = 20, P21: n = 20, P30: n =9. *p<0.05.

## Results

### Ontogeny of Persistent (9-day) Auditory Fear Memory

We first explored the ontogeny of persistent auditory fear memory across the two C57BL/6 substrains. We chose a 9-day retrieval to match the timeline of Gogolla et al. (2009)^56^, a reference in the field using C57BL/6J mice, and that of our extinction experiments. We first explored the ontogeny of 9-day auditory fear memory in Jax mice to later compare with Taconic mice. We trained P16, P18, P21 and P30 Jax mice in auditory fear conditioning and tested for auditory memory retrieval 9 days later. Jax mice of all age groups successfully acquired the tone-shock fear association, with P21 Jax mice showing increased freezing compared to P30 mice during tones 1-3, and P16 Jax mice showing increased freezing at tone 4 compared to P18 and P30 (**Figure 1B**: RM Two-way ANOVA: F_interaction 13, 212_ = 2.31, *p* = 0.0071; F_tone 4.3, 212_ = 34.68, *p* < 0.0001; F_age 3, 49_ = 3.37, *p* = 0.026. Tukey’s post-hoc, Tone 1: P21 vs. P30 *p* = 0.0094; Tone 2: P21 vs. P30 *p* = 0.022; Tone 3: P21 vs P30 *p* = 0.011; Tone 4: P16 vs. P18 *p* = 0.03, P16 vs. P30 *p* = 0.013). At the 9-day retrieval test, P16 and P18 Jax mice showed reduced freezing compared to P21 and P30 mice (**Figure 1C**: one-way ANOVA: F_3, 49_ = 16.42, *p* < 0.0001; Tukey’s post hoc test: P16 vs P21: *p* = 0.0002; P16 vs P30: *p <* 0.0001; P18 vs. P21: *p* = 0.0075; P18 vs. P30: *p* < 0.0001). This suggests that Jax mice first show 9-day persistent auditory fear memory at P21.

We then conducted similar experiments in Taconic mice at the same ages. Taconic mice successfully acquired the fear association, but P30 mice showed reduced freezing at tones 1, 2 and P16 mice showed increased freezing at tone 6 compared to P18 and P21 (**Figure 1D**: RM Two-way ANOVA: F_interaction 14, 244_ = 2.44, *p* = 0.0032; F_tone 4.60, 244_ = 41.03, *p* < 0.0001; F_age 3, 53_ = 1.3, *p* = 0.283. Tukey’s post-hoc, Tone 1: P18 vs. P30 *p* = 0.03, P21 vs. P30 *p* = 0.022; Tone 2: P16 vs. P30 *p* = 0.023, P18 vs. P30 *p* = 0.0001, P21 vs. P30 *p* = 0.0001; Tone 6: P16 vs. P18 *p* = 0.0009, P16 vs. P21 p=0.023). During the 9-day auditory fear retrieval test, Taconic mice of all ages showed equivalent freezing (**Figure 1E**: one-way ANOVA: F_3, 52_ = 3.83, p = 0.015), indicating successful memory retrieval from the earliest age tested. Direct comparison of freezing between substrains at each age also shows increased freezing in Taconic mice at P16 and P18 compared to Jax mice (**Figure S1A-E**: CS1-2: unpaired t-test, P16: t_16_ = 3.775, *p* = 0.0017; P18: t_32_ = 2.798, *p* = 0.0086; including for CS1-5: **Figure S1F-I**: CS1-5: unpaired t-test, P16: t_16_ = 3.715, *p* = 0.0019; P18: t_32_ = 2.316, *p* = 0.0271), suggesting that C57BL/6NTac Taconic mice display an earlier onset of persistent memory compared to Jax mice, starting at P16.

### Ontogeny of C57BL/6J (Jax) spontaneous recovery and renewal

To determine whether the developmental emergence of adult-type fear extinction differs among C57BL/6 substrains, we next explored the ontogeny of spontaneous recovery and renewal following auditory fear extinction in Jax and Taconic mice. Given that Jax mice show infantile amnesia for 9-day auditory memory prior to P21 (**Figure 1**), to exclude the effects of forgetting we fear trained Jax mice starting at P21, including P25 and P30. During fear acquisition, Jax male mice of all ages showed incremental freezing across tones, suggesting they acquired the tone-shock association (**Figure 2B**: repeated measures (RM) two-way ANOVA: F_interaction 8.6, 138.1_ = 2.14, *p* = 0.032; F_tone 4.3, 138.1_ = 38.98, *p* < 0.0001; F_age 2, 32_ = 7.48, *p* = 0.0022). Jax P21 males exhibited enhanced freezing during the first three tones of fear acquisition (**Figure 2B**: Tukey’s post-hoc test Tone 1: P21 vs. P25 *p* = 0.0095, P21 vs. P30 *p* = 0.0039; Tone 2: P21 vs. P25 *p* = 0.015; Tone 3: P21 vs P25 *p* = 0.0058). Female Jax mice also showed increased freezing across the training session, with P21 females displaying enhanced freezing during tone 2 compared to P25 and P30 females (**Figure 2I**: RM Two-way ANOVA: F_interaction 6.98, 97.7_ = 3.90, *p* = 0.0009; F_tone 3.5, 97.7_ = 68.91, *p* < 0.0001; F_age 2, 28_ = 3.17, *p* = 0.057. Tukey’s post-hoc, Tone 2: P21 vs. P25 *p* < 0.0001, P21 vs. P30 *p* = 0.037).

**Figure 2.**
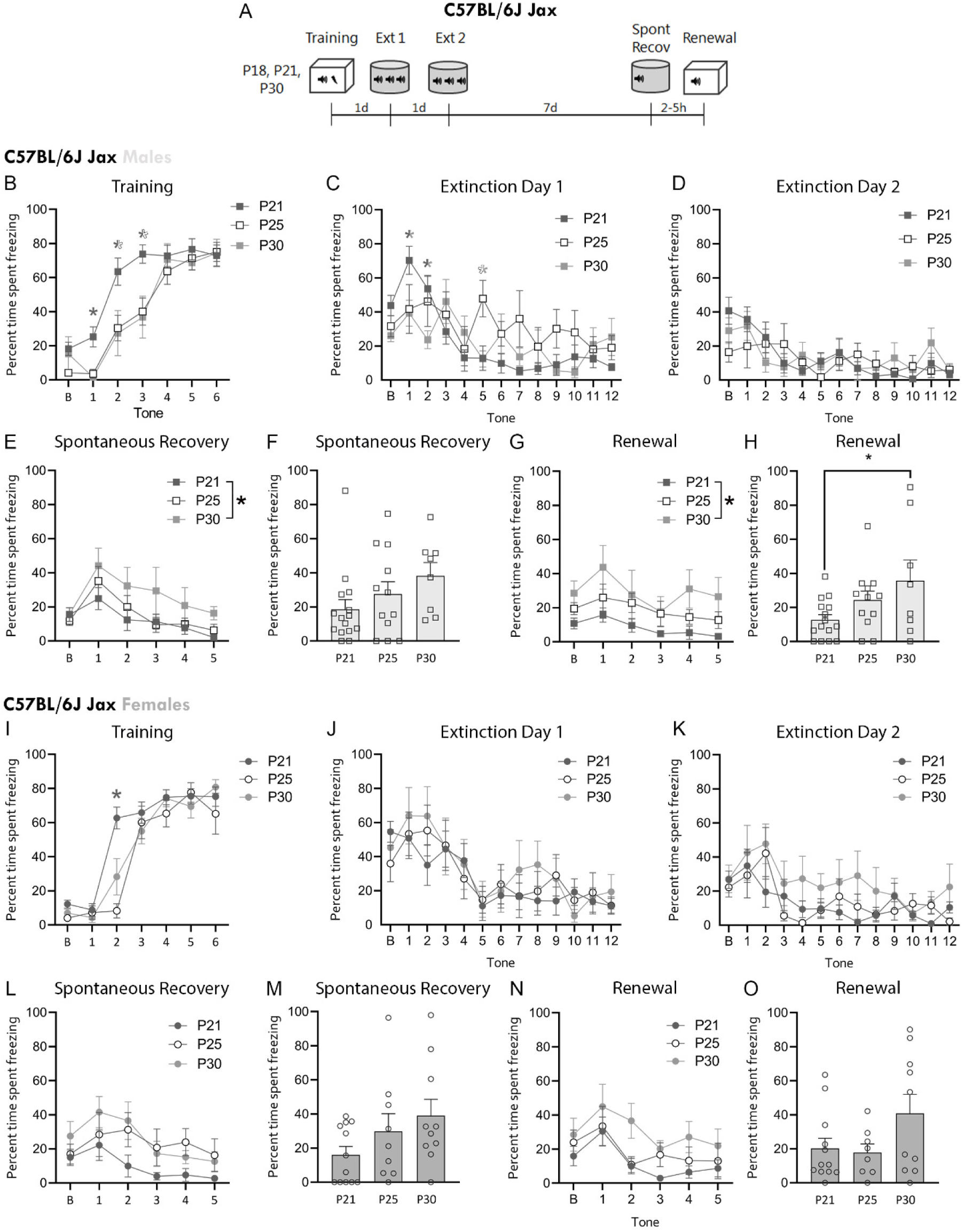
C57BL/6J (Jax) mice show spontaneous recovery by P21, but males show increased renewal by P30. **A.** Schematic of fear extinction design. **B-H.** Male Jax data. Tone-by-tone freezing during fear training (**B**;* represents a difference between P21 vs. P25 and P30 at tone 1, represents a difference between P21 vs P25 at tones 2 and 3), extinction day 1(**C; \*** represents a difference between P21 vs 30 at tones 1, 2; and *between P25 vs P30 at tone 5), extinction day 2 (**D**) and spontaneous recovery (**E**)**. F.** Average freezing during CS 1-2 of spontaneous recovery. Tone-by tone (**G**) and average freezing (**H**) at renewal**. I-O.** Female Jax mice data. Tone-by-tone freezing during fear training (**I**), extinction day 1 (**J**), extinction day 2 (**K**) and spontaneous recovery (**L**)**. M.** Average freezing during CS 1-2 of spontaneous recovery. Tone-by tone (**N**; CS 1-2) and average freezing (**O**) at renewal. Males: P21: *n* = 15, P25: *n* = 12, P30: *n* = 8. Females: P21: *n* = 12, P25: *n* = 9, P30: *n* = 10. *p<0.05.

On extinction day 1, all Jax male groups displayed high levels of freezing during the initial tone presentations, with P21 males showing higher freezing relative to P30 males during tones 1 and 2, and P25 males freezing more than P30 during tone 5 (**Figure 1C**: RM Two-way ANOVA: F_interaction 15.2, 242_ = 1.94, *p* = 0.0197; F_tone 7.59, 242_ = 10.01, *p* < 0.0001; F_age 2, 32_ = 2.36, *p* = 0.11. Tukey’s post-hoc Tone 1: P21 vs. P30 *p* = 0.031, Tone 2: P21 vs. P30 *p* = 0.013, Tone 5: P25 vs. P30 *p* = 0.025). In contrast, female Jax mice showed no age-dependent differences on extinction day 1 (**Figure 1J**: RM Two-way ANOVA: F_interaction 13, 184_ = 0.33, *p* = 0.99; F_tone 6.6, 184.2_ = 7.20, *p* < 0.0001; F_age 2, 28_ = 0.70, *p* = 0.50). On extinction day 2, both Jax males and females showed equivalent levels of freezing across the session (**Figure 1D** males: RM Two-way ANOVA F_interaction 12.2, 194.9_ = 0.92, *p* = 0.527; F_tone 6.1, 194.9_ = 4.83, *p* = 0.0001; F_age 2, 32_ = 0.126, *p* = 0.882; **Figure 1K** females: RM Two-way ANOVA F_interaction 11.35, 158.9_ = 1.05, *p* = 0.40; F_tone 5.67, 158.9_ = 6.47, *p* < 0.0001; F_age 2, 28_ = 2.73, *p* = 0.083). Overall, all Jax mice were able to acquire and extinguish the tone-shock association.

We then assessed spontaneous recovery a week following extinction learning. Jax males show similar average freezing during spontaneous recovery across ages, but tone-by tone comparisons shows slightly higher freezing in P30 mice compared to P21 (**Figure 2E**: RM Two-way ANOVA: F_interaction 6.5, 104.5_ = 0.294, *p* = 0.948; F_tone 3.3, 104.5_ = 9.91, *p* < 0.0001; F_age 2, 32_ = 3.63, *p* = 0.0379; Tukey’s post-hoc, P21 vs 30 p=0.0301; **Figure 2F**: One-way ANOVA: F_2, 32_ = 1.92, *p* = 0.163). Female Jax mice display equivalent spontaneous recovery across ages (**Figure 2L**: RM Two-way ANOVA: F_interaction 6.2, 87.4_ = 0.702, *p* = 0.655; F_tone 3.1, 87.4_ = 6.05, *p* = 0.0007; F_age 2, 28_ = 2.95, *p* = 0.069; **Figure 2M**: One-way ANOVA: F_2,28_ = 2.18, *p* = 0.131), indicating that Jax male and female mice show similar spontaneous recovery from as early as P21. However, when tested for fear renewal, P30 Jax males showed higher freezing compared to P21 (**Figure 2G**: RM Two-way ANOVA: F_interaction 6.5, 104.8_ = 0.42, *p* = 0.876; F_tone 3.3, 104.8_ = 3.64, *p* = 0.013; F_age 2, 32_ = 6.31, *p* = 0.0049; post-hoc P21 vs P30 p=0.0040; **Figure 2H**: One-way ANOVA: F_2, 32_ = 3.34, p = 0.048; Tukey’s post-hoc test: P21 vs. P30 *p* = 0.042). Jax females showed comparable renewal across the tested ages (**Figure 2N**: RM Two-way ANOVA: F_interaction 5.5, 77.2_ = 0.481, *p* = 0.81; F_tone 2.8, 77.2_ = 6.69, *p* = 0.0006; F_age 2, 28_ = 3.15, *p* = 0.059; **Figure 2O**: One-way ANOVA: F_2, 27_ = 2.45, *p* = 0.11). These data suggest that while the onset of spontaneous recovery occurs at P21 in Jax male and female mice, Jax male mice display increased renewal at P30.

### Ontogeny of C57BL/6NTac (Taconic) spontaneous recovery and renewal

To explore the developmental timeline of spontaneous recovery and renewal in C57BL/6NTac Taconic mice, we chose slightly different ages based on their early emergence of remote fear (**Figure 1**). We fear trained Taconic mice at P18, 21, and 30. Taconic male mice showed comparable levels of freezing at fear acquisition across ages (**Figure 3B**: RM Two-way ANOVA, males: F_interaction 7.6, 121_ = 1.33, *p* = 0.24; F_tone 3.78, 121_ = 19.85, *p* < 0.0001; F_age 2, 32_ = 2.27, *p* = 0.120). In contrast, Taconic female P30 mice showed decreased freezing compared to P18 and P21 at fear acquisition (**Figure 3I**: RM Two-way ANOVA, females: F_interaction 8.1, 125_ = 1.33, *p* = 0.23; F_tone 4.03, 124.9_ = 28.35, *p* < 0.0001; F_age 2, 31_ = 7.63, *p* = 0.002; P18 vs P30 p=0.0015, P21 vs P30 p=0.0310). During extinction day 1, P30 Taconic males showed higher freezing compared to P18 (**Figure 3C**: RM Two-way ANOVA, males: F_interaction 14, 257_ = 0.801, *p* = 0.67; F_tone 7.1, 257_ = 9.82, *p* < 0.0001; F_age 2, 36_ = 3.64, *p* = 0.036, P18 vs P30 p=0.028), but females showed no age-dependent differences (**Figure 3J**: females: F_interaction 15, 259_ = 1.20, *p* = 0.27; F_tone 7.62, 259_ = 7.11, *p* < 0.0001; F_age 2, 34_ = 3.51, *p* = 0.041; post-hocs p>0.055). Neither sex showed age-related differences during extinction day 2 (**Figure 3D**: RM Two-way ANOVA, males: F_interaction 14.3, 257_ = 0.513, *p* = 0.927; F_tone 7.14, 257_ = 6.70, *p* < 0.0001; F_age 2, 36_ = 2.05, *p* = 0.143. **Figure 3K**, RM Two-way ANOVA, females: F_interaction 14, 239_ = 1.31, *p* = 0.203; F_tone 7, 239_ = 8.79, *p* < 0.0001; F_age 2, 24_ = 2.99, *p* = 0.064). These data suggest sex- and age-specific differences in fear acquisition and extinction across C57BL/6 substrains, particularly in P30 mice.

**Figure 3.**
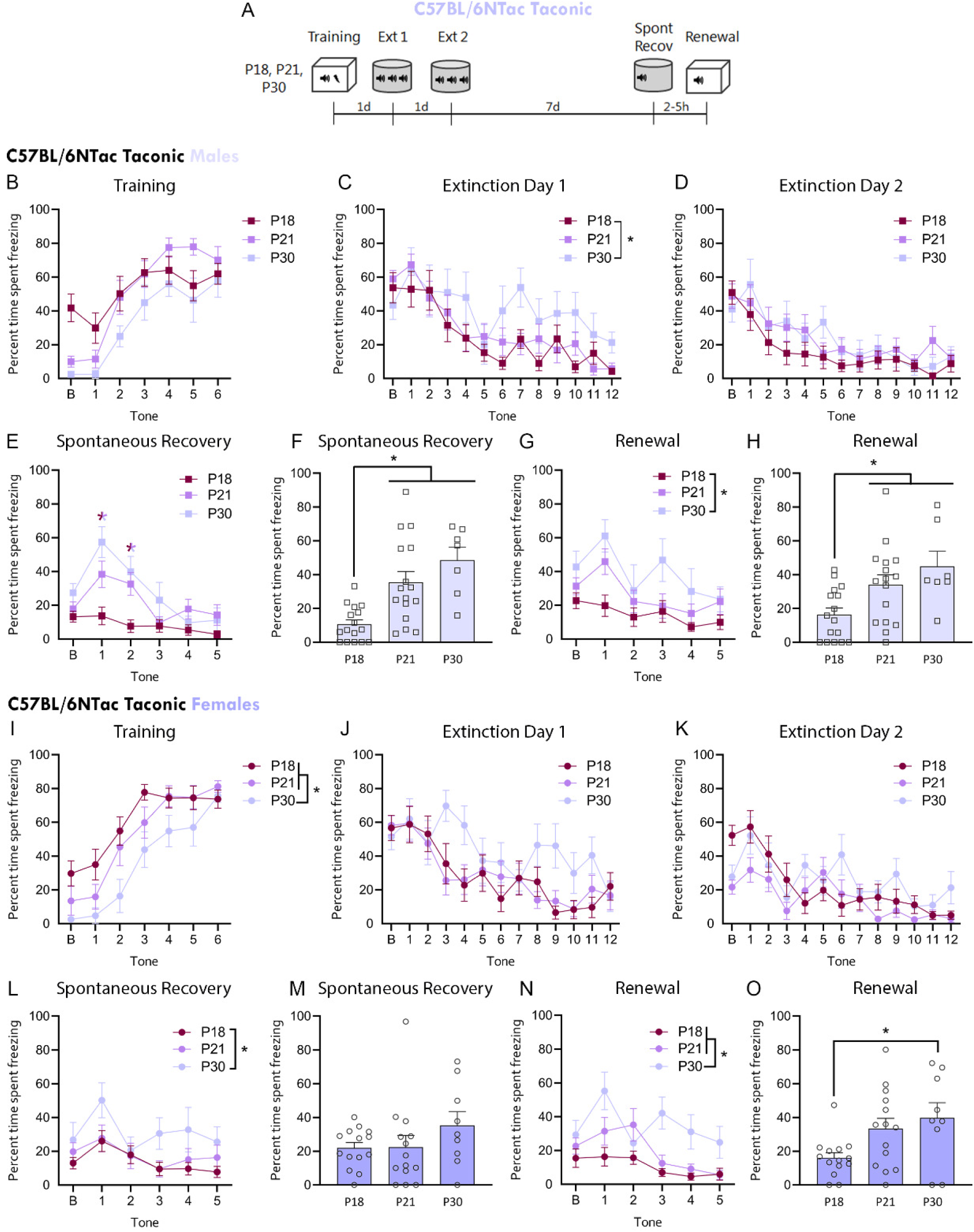
Developmental profile of fear training and extinction in C57BL/6NTac (Taconic) mice. **A.** Schematic of fear extinction design. **B-H.** Male Taconics data. Tone-by-tone freezing during fear training (**B**), extinction day 1(**C**), extinction day 2 (**D**) and spontaneous recovery (**E**; * represents a difference between P18 vs 21 and 30 at tones 1 and 2). **F.** Average freezing during CS 1-2 of spontaneous recovery. Tone-by tone (**G**) and average freezing (**H**) at renewal**. I-O.** Female Taconics data. Tone-by-tone freezing during fear training (**I**), extinction day 1 (**J**), extinction day 2 (**K**) and spontaneous recovery (**L**)**. M.** Average freezing during CS 1-2 of spontaneous recovery. Tone-by tone (**N**; CS 1-2) and average freezing (**O**) at renewal. Males: P18: *n* = 16, P21: *n* = 16, P30: *n* = 7. Females: P18: *n* = 14, P21: *n* = 14, P30: *n* = 9. *p<0.05.

One week following extinction learning, mice were tested for spontaneous recovery. P18 Taconic males showed low freezing levels compared to P21 and P30 male mice at spontaneous recovery (**Figure 3E**: RM Two-way ANOVA: F_interaction 6.7, 120_ = 2.49, *p* = 0.022; F_tone 3.34, 120_ = 12.8, *p* < 0.0001; F_age 2, 36_ = 10.28, *p* = 0.0003; Tukey’s post-hoc - Tone 1: P18 vs. P21 *p* = 0.037, P18 vs. P30 *p* = 0.0049; Tone 2: P18 vs. P21 *p* = 0.0093, P18 vs. P30 *p* = 0.028; **Figure 3F**: One-way ANOVA: F_2, 36_ = 11.59, *p* = 0.0001; Tukey’s post-hoc: P18 vs. P21 *p* = 0.0024, P18 vs. P30 *p* = 0.0003), consistent with an absence of spontaneous recovery in P18 Taconic males. In contrast, Taconic females show a largely similar profile across ages for spontaneous recovery, with the exception of increased freezing by P30 Taconic females in the tone-by-tone (but not average freezing) comparison (**Figure 3L**: RM Two-way ANOVA: F_interaction 7.4, 125.7_ = 0.712, *p* = 0.67; F_tone 3.7, 125.7_ = 5.13, *p* = 0.001; F_age 2, 34_ = 3.57, *p* = 0.039, P18 vs P30 p=0.0377; **Figure 3M**: One-way ANOVA: F_2, 34_ = 1.33, *p* = 0.278). This points to an earlier onset of spontaneous recovery in Taconic females (P18) compared to Taconic males (P21).

During the fear renewal test, both Taconic male and female mice showed higher freezing at older ages, starting at P21 for males and P30 for females (**Figure 3G**: Males: RM Two-way ANOVA, F_interaction 6.6, 118.6_ = 1.14, *p* = 0.345; F_tone 3.3, 118.6_ = 7.01, *p* = 0.0001; F_age 2, 36_ = 5.62, *p* = 0.0075; Tukey’s post-hoc, P18 vs. P30 *p* = 0.0068. **Figure 3H**: One-way ANOVA, F_2, 36_ = 5.64, *p* = 0.0074; Tukey’s post-hoc test: P18 vs. P21 *p* = 0.0498, P18 vs. P30 *p* = 0.011. **Figure 3N**: Females: RM Two-way ANOVA: F_interaction 6.2, 106.6_ = 1.93, *p* = 0.08; F_tone 3.1, 106.6_ = 6.11, *p* = 0.0006; F_age 2, 34_ = 12.28, *p* < 0.0001; Tukey’s post-hoc, P18 vs. P30 *p* <0.0001; P21 vs. P30 *p* = 0.0072. **Figure 3O**: One-way ANOVA: F_2, 34_ = 4.30, *p* = 0.022; Tukey’s post-hoc test: P18 vs. P30 *p* = 0.028), suggesting that the onset of renewal occurs around P21-P30 in C57BL/6NTac mice.

Together, these data point to sex and substrain differences in fear processing. Taconic female mice display earlier spontaneous recovery (from P18) compared to Taconic male, Jax male and female mice. Additional between-strain analyses confirmed no strain or sex differences in spontaneous recovery at P21 and P30 (**Supplementary Figure 2B-C**: P21 - Two-way ANOVA: F_Interaction 1, 51_ = 1.189, *p* = 0.281; F_Strain 1, 51_ = 2.546, *p* = 0.117; F_Sex 1, 51_ = 0.963, *p* = 0.331. P30 - F_Interaction 1, 30_ = 0.670, *p* = 0.420; F_Strain 1, 30_ = 0.145, *p* = 0.706; F_Sex 1, 30_ = 0.521, *p* = 0.480). Taconic males show earlier increases in fear renewal (P21) compared to Taconic females and Jax males (both at P30), matching the timing of Jax females, who show stable levels of renewal from P21. This was reinforced by between-strain analyses of these data, which showed increased freezing by P21 Taconic males compared to P21 Jax males (**Supplementary Figure 2D-E**: F_Interaction 1, 53_ = 0.608, *p* = 0.439; F_Strain 1, 53_ = 10.27, *p* = 0.0023, F_Sex 1, 53_ = 0.399, *p* = 0.530. Tukey’s post-hoc test, Taconic males vs Jax males *p* = 0.0093), and no substrain differences by P30 (F_Interaction 1, 30_ = 0.231, *p* = 0.635; F_Strain 1, 30_ = 0.147, *p* = 0.704, F_Sex 1, 30_ = 0.0130, *p* = 0.997), suggesting comparable levels of renewal by adolescence across sexes and substrains.

## Discussion

Here we compared the ontogeny of 9-day persistent auditory fear memory, spontaneous recovery and renewal between two C57BL/6 mouse substrains known to display genotypic and phenotypic differences in fear behavior in adulthood^19,36^.

C57BL/6NTac Taconic mice show an early emergence of adult-like fear behavior, including an earlier onset of 9-day memory persistence, as well as earlier spontaneous recovery in females compared to Jax mice. Taconic male mice also show earlier renewal compared to Jax. These findings uncover further divergence in fear behaviors among seemingly equivalent mouse substrains, and demonstrate that substrain differences may emerge quite early in development, carrying important implications for experimental design and interpretation.

We found that C57BL/6J Jax mice display slightly longer infantile amnesia compared to Taconic mice, with 9-day memory persistence only emerging by P21 in Jax compared to P16 in Taconic mice. Interestingly, our previous work showed that C57BL/6J Jax mice were unable to retrieve remote 30-day auditory fear memory when trained at P21 but not P25^50^). This differential developmental timeline for 9-day and 30-day old memories parallels evidence of distinct neural requirements for memory retrieval at varying intervals post-encoding^62,63^, suggesting the existence of a temporal gradient in the ontogeny of memory persistence, with successful retrieval of older (30-day) memories emerging later in development compared to that of remote memories with a shorter retention interval (9-day). Importantly, given the increased reliance on cortical networks at longer retention intervals^58–61,64^, and the known delayed maturation of the medial prefrontal cortex (mPFC)^65,66^, cortical maturation may be an important factor in guiding this differential timing within the ontogenesis of memory persistence.

The developmental ages assessed in our experiments coincide with synaptic and anatomical changes in brain regions such as amygdala, mPFC and hippocampus^67–80^, which are implicated in adult fear acquisition, extinction^76,81–85^ and renewal^86–88^. This raises the intriguing possibility that the reported behavioral differences may be accompanied by substrain and/or sex differences in the maturation of corticolimbic circuitry. Interestingly, individual differences in amygdala-prefrontal cortex functional connectivity in humans are present from childhood^89–91^, with known contributions of genetic variations to amygdala activity, amygdala-cortical connectivity and emotional reactivity in children and adolescents^91^. Importantly, even though extrinsic modulators such as stress alter the developmental timeline of fear-related brain pathways in humans^89,90^ and rodents^92,93^, the exact impact of intrinsic factors such as genetic differences on the maturation of emotional processing and its neural circuitry remains underexplored.

While we noted some sex and substrain differences in both fear and extinction learning, all our age groups successfully extinguished fear. This contrasts with several studies reporting deficits in fear extinction in adolescent rats and mice^40,94–99^. The absence of an adolescent extinction deficit in our data may be attributed to differences in fear extinction protocols (for detailed discussion, see ^100^) , as a few studies found that doubling or extending the extinction protocol (as was done here) can rescue the extinction deficit in adolescent mice and rats^102–104^. Furthermore, although our analyses concentrated on freezing behavior, other fear behaviors such as rearing, pivoting, and grooming vary by age^105^, suggesting that additional developmental substrain differences may be uncovered by an expanded behavioral analysis.

Our data uncovered sex differences in fear processing in early life, particularly in the C57BL/6NTac substrain. Although research on the intersection between strain and sex differences is sparse, some work points to interactions between strain and sex in mouse social, fear, and anxiety-like behaviors in adulthood^106–108^. In development, Park et al. (2017) reported sex differences in fear behavior in rats, with female rats displaying earlier renewal and 5-day spontaneous recovery compared to male rats assessed at P18 (but see ^109^). While our data was not as definitive, Jax males did display a trend toward a slightly delayed profile of spontaneous recovery and renewal compared to females (**Figure 2**), and female Taconic mice had an earlier plateauing of spontaneous recovery compared to Taconic males (**Figure 3**). These findings converge on an overall earlier maturation profile for female rodent fear behavior, but additional experiments are necessary to ascertain the robustness of C56BL/6 mouse sex differences in developmental fear. Another important variable when interpreting sex differences in early life is puberty, especially since pubertal onset itself differs between rodent strains^110–112^. Given that the onset of puberty can impact cortical maturation^113^, and that gonadal hormones affect the neural correlates of fear learning^114–124^, substrain differences in pubertal onset and/or in hormone levels prior to pubertal onset, particularly in female mice, may affect the ontogeny of fear processing.

Overall, this work suggests that C57BL/6NTac mice reach a mature, adult-like fear profile earlier than Jax mice. These substrain differences in the ontogenesis of fear processing emphasize the importance of the choice of rodent strain and substrain in any experimental design and urge careful interpretation of findings related to rodent fear learning and extinction across ages. While we only examined two C57BL/6 substrains, the known strain differences in adult fear expression^125^ suggest more widespread variance among the ontogenetic profiles of fear processing across mouse strains and substrains. This highlights the need for increased transparency when reporting strain/substrain information, particularly when there are difficulties replicating data.

Critically, substrain variations in the ontogenetic profile of fear behavior can also be leveraged toward experiments probing the link between brain maturation and behavioral emergence, offering unique insight into the neural basis of behavior. Together, our findings provide a base for future experiments exploring the developmental profile of strain and substrain differences, and highlight how these can serve as powerful tools to study the effects of individual and genetic differences on phenotypic behaviors.

## Acknowledgements

This work was supported by grants from the Natural Science and Engineering Council of Canada (RGPIN-2017-06344) to MA-C. We thank Bhairavei Gnanamanogaran, Mehreen Inayat, and Unza Mumtaz for general support with animal research and colony management in our lab.

## Author contributions

HP, KN performed research. HP, KN analyzed data. HP and MA-C designed research and wrote the paper.

## Supplemental information

Document S1: Supplemental Figures S1–S2, Supplemental Statistical Analysis Table S1.

---

**Supplemental Figure S1.**
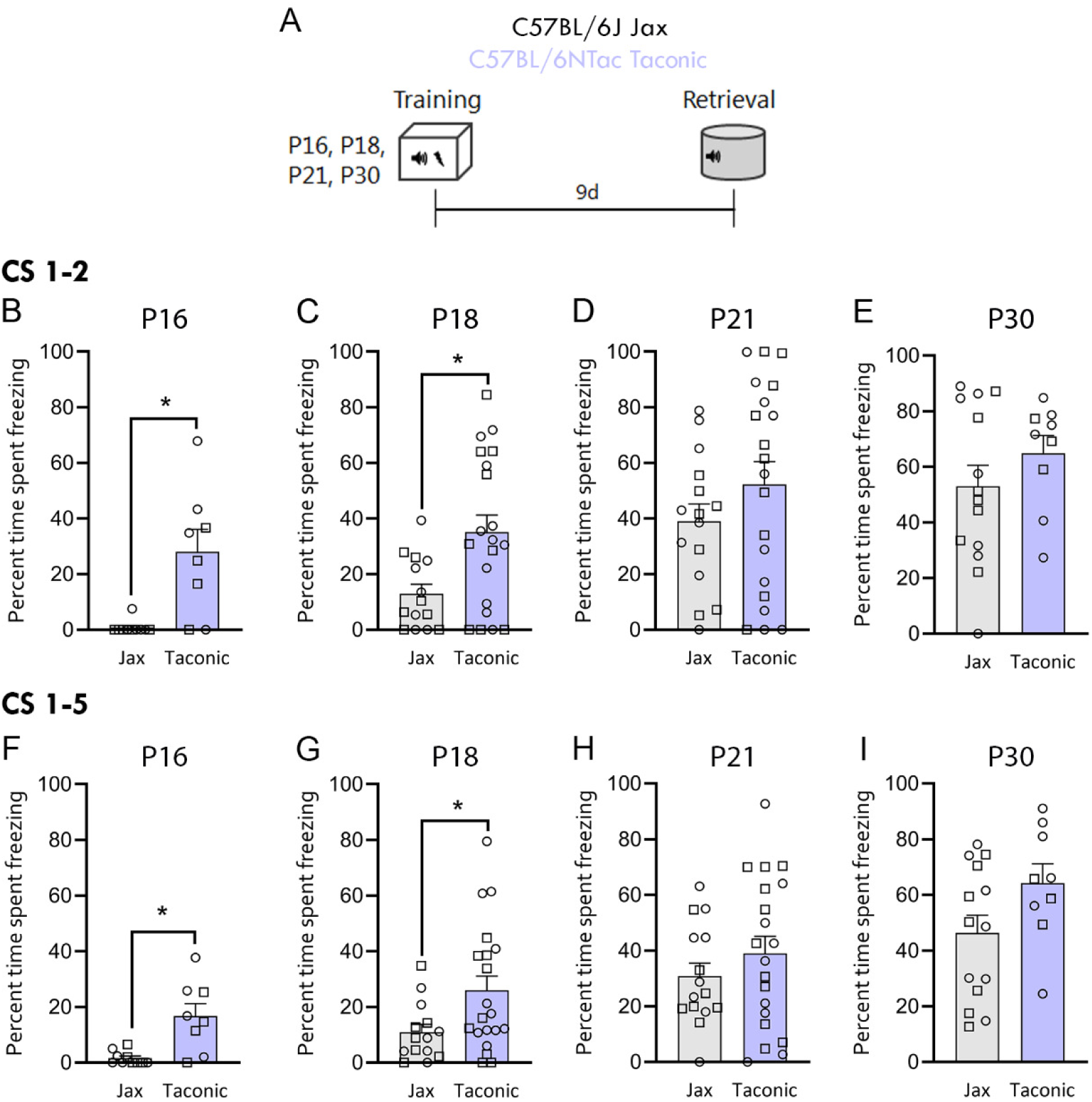
Between-strain comparison of the ontogeny of persistent auditory fear memory in C57BL/6J (Jax) and C57BL/6NTac (Taconic) mice. These are the same data displayed in **Figure 1**, only here showing the comparison between substrains. **A.** Schematic of behavioral design. Mice were fear trained on day 1 and tested for 9d retrieval on day 10. **B-C.** Taconic mice show increased freezing during retrieval testing at P16 and P18 compared to Jax mice (average freezing to CS1-2: unpaired t-test, P16: t_16_ = 3.775, *p* = 0.0017; P18: t_32_ = 2.798, *p* = 0.0086; P21: t_33_ = 1.208, *p* = 0.2356; P30: t_21_ = 1.100, *p* = 0.2839). **D-E.** Similar results are seen when averaging all 5 tones (CS 1-5) of the retrieval test, with Taconic mice showing increased freezing at P16 compared to Jax (unpaired t-test, P16: t_16_ = 3.715, *p* = 0.0019; P18: t_32_ = 2.316, *p* = 0.0271; P21: t_33_ = 1.010, *p* = 0.3198; P30: t_21_ = 1.854, *p* = 0.0779). P16 Taconic: n = 8, P18 Taconic: n = 20, P21 Taconic: n = 20, P30 Taconic: n =9. P16 Jax: n = 10, P18 Jax: n =14, P21 Jax: n = 15, P30 Jax: n =14. *p<0.05.

**Supplemental Figure S2.**
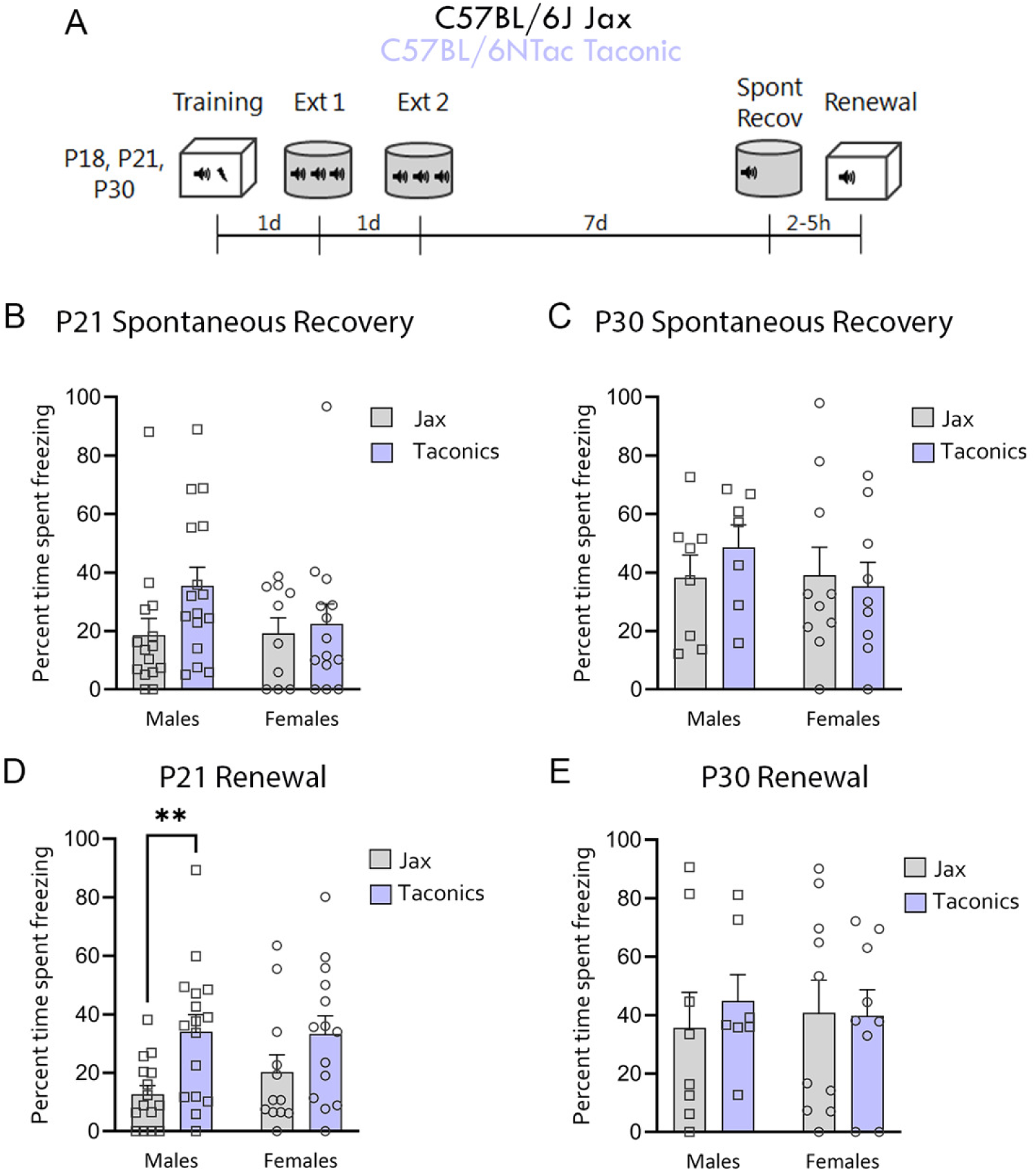
Between-strain comparison of spontaneous recovery in C57BL/6NTac (Taconic) and C57BL/6J (Jax) mice. These are the same data displayed in **Figures 2 and 3**, only here showing the comparison between substrains. **A.** Experimental design. **B-C.** Spontaneous recovery at P21 (**B;** CS1-2: Two-way ANOVA: F_Strain 1, 51_ = 2.546, *p* = 0.117; F_Sex 1, 51_ = 0.963, *p* = 0.331; F_Interaction 1, 51_ = 1.189, *p* = 0.281) and P30 (**C;** CS1-2: Two-way ANOVA: F_Strain 1, 27_ = 0.416, *p* = 0.524; F_Sex 1, 27_ = 0.992, *p* = 0.328; F_Interaction 1, 27_ = 0.348, *p* = 0.560) showed no differences between substrains. **D-E.** Renewal. At P21 (**D**), Jax male mice demonstrate a lack of renewal compared to Taconic mice (F_Strain 1, 53_ = 10.27, *p* = 0.0023, F_Sex 1, 53_ = 0.399, *p* = 0.530, F_Interaction 1, 53_ = 0.608, *p* = 0.439; Tukey’s post-hoc, Taconic males vs. Jax males *p* = 0.024, Taconic females vs. Jax males p = 0.039), an effect absent at P30 (**E**; F_Strain 1, 27_ = 0.108, *p* = 0.746, F_Sex 1, 27_ = 0.002, *p* = 0.964, F_Interaction 1, 27_ = 0.256, *p* = 0.617). Jax males P21: *n* = 15,. Jax females P21: *n* = 12. Taconic males: P21: *n* = 16, Taconic females P21: *n* = 14. Jax males P30: *n* = 8, Jax females P30: *n* = 7. Taconic males P30: *n* = 7, Taconic females P30: *n* = 9. ** p<0.005.

**Supplemental Table 1.**
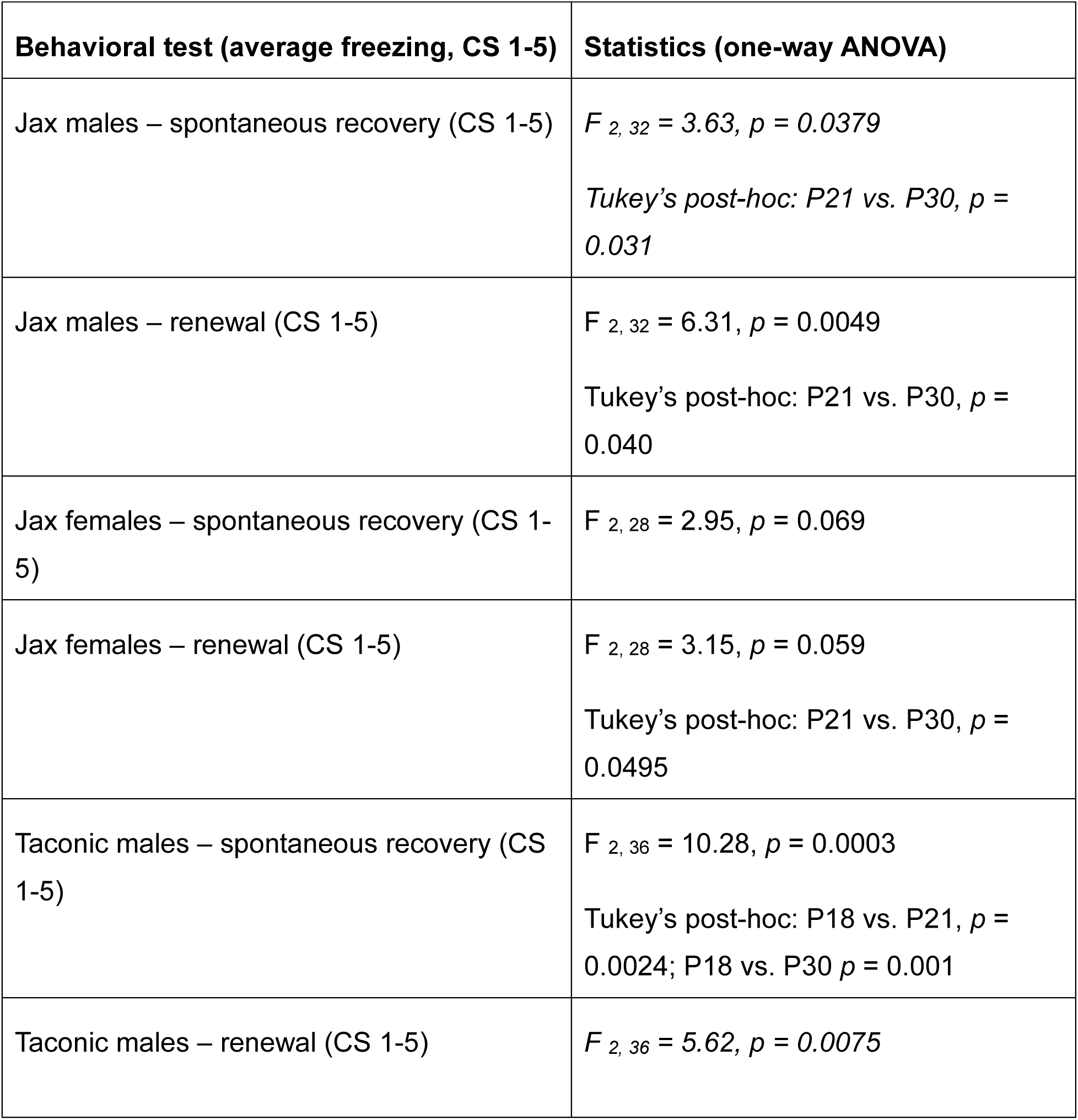
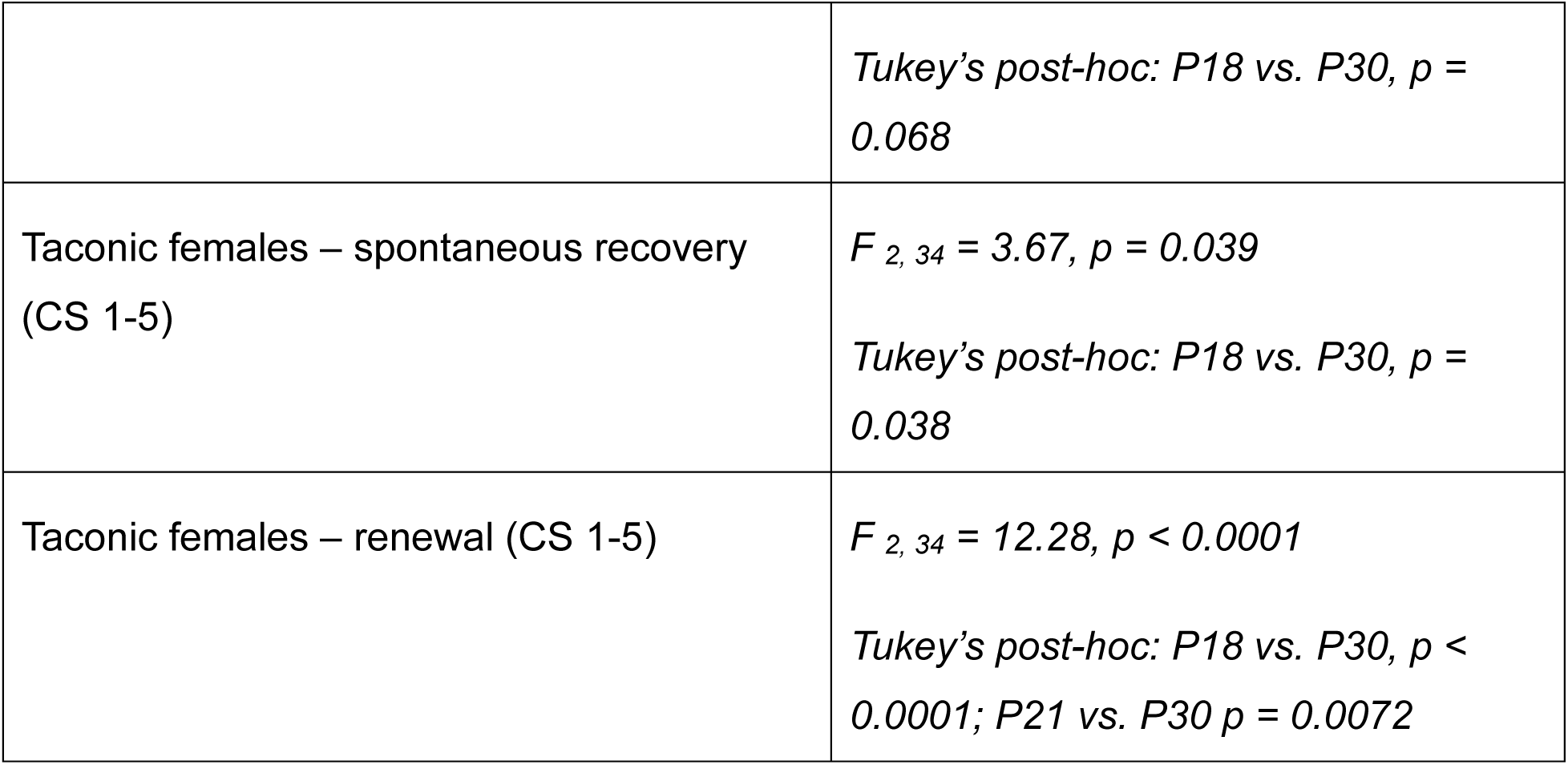
Statistical analysis of freezing during CS1-5 at test.

| Behavioral test (average freezing, CS 1-5) | Statistics (one-way ANOVA) |
| --- | --- |
| Jax males – spontaneous recovery (CS 1-5) | $F_{2, 32} = 3.63$, $p = 0.0379$<br>Tukey's post-hoc: P21 vs. P30, $p = 0.031$ |
| Jax males – renewal (CS 1-5) | $F_{2, 32} = 6.31$, $p = 0.0049$<br>Tukey's post-hoc: P21 vs. P30, $p = 0.040$ |
| Jax females – spontaneous recovery (CS 1-5) | $F_{2, 28} = 2.95$, $p = 0.069$ |
| Jax females – renewal (CS 1-5) | $F_{2, 28} = 3.15$, $p = 0.059$<br>Tukey's post-hoc: P21 vs. P30, $p = 0.0495$ |
| Taconic males – spontaneous recovery (CS 1-5) | $F_{2, 36} = 10.28$, $p = 0.0003$<br>Tukey's post-hoc: P18 vs. P21, $p = 0.0024$; P18 vs. P30 $p = 0.001$ |
| Taconic males – renewal (CS 1-5) | $F_{2, 36} = 5.62$, $p = 0.0075$ |
|  | *Tukey's post-hoc: P18 vs. P30, $p = 0.068$* |
| Taconic females – spontaneous recovery (CS 1-5) | $F_{2, 34} = 3.67, p = 0.039$<br>*Tukey's post-hoc: P18 vs. P30, $p = 0.038$* |
| Taconic females – renewal (CS 1-5) | $F_{2, 34} = 12.28, p < 0.0001$<br>*Tukey's post-hoc: P18 vs. P30, $p < 0.0001$; P21 vs. P30 $p = 0.0072$* |

## References

1. Hartley, C. A. & Casey, B. J. Risk for anxiety and implications for treatment: Developmental, environmental, and genetic factors governing fear regulation. Ann N Y Acad Sci 1304, (2013).

2. King, G., Graham, B. M. & Richardson, R. Individual differences in fear relapse. Behaviour Research and Therapy 100, (2018).

3. Hettema, J. M., Annas, P., Neale, M. C., Kendler, K. S. & Fredrikson, M. A twin study of the genetics of fear conditioning. Arch Gen Psychiatry 60, (2003).

4. Purves, K. L. et al. Evidence for distinct genetic and environmental influences on fear acquisition and extinction. Psychol Med 53, (2023).

5. Zhylin, M., Mendelo, V., Bondarevych, S., Kokorina, Y. & Tatianchykov, A. Genetic Basis of Emotional Regulation: Integrative Analysis of Behavioral and Neurobiological Data. OBM Neurobiol 08, 1–21 (2024).

6. Kastrati, G. et al. Genetic influences on central and peripheral nervous system activity during fear conditioning. Transl Psychiatry 12, (2022).

7. Bush, D. E. A., Sotres-Bayon, F. & LeDoux, J. E. Individual differences in fear: Isolating fear reactivity and fear recovery phenotypes. in *Journal of Traumatic Stress* vol. 20 (2007).

8. Zeidan, M. A. et al. Test-Retest Reliability during Fear Acquisition and Fear Extinction in Humans. CNS Neurosci Ther 18, (2012).

9. MacPherson, K. et al. Temporal factors in the extinction of fear in inbred mouse strains differing in extinction efficacy. Biol Mood Anxiety Disord 3, (2013).

10. Caspi, A., Hariri, A. R., Holmes, A., Uher, R. & Moffitt, T. E. Genetic Sensitivity to the Environment: The Case of the Serotonin Transporter Gene and Its Implications for Studying Complex Diseases and Traits. Focus (Madison) 8, (2010).

11. Skelton, K., Ressler, K. J., Norrholm, S. D., Jovanovic, T. & Bradley-Davino, B. PTSD and gene variants: New pathways and new thinking. Neuropharmacology vol. 62 Preprint at 10.1016/j.neuropharm.2011.02.013 (2012).

12. Caspi, A. & Moffitt, T. E. Gene-environment interactions in psychiatry: Joining forces with neuroscience. Nature Reviews Neuroscience vol. 7 Preprint at 10.1038/nrn1925 (2006).

13. Caspi, A., Hariri, A. R., Holmes, A., Uher, R. & Moffitt, T. E. Genetic Sensitivity to the Environment: The Case of the Serotonin Transporter Gene and Its Implications for Studying Complex Diseases and Traits. Focus (Madison) 8, (2010).

14. Rutter, M. Biological implications of gene-environment interaction. J Abnorm Child Psychol 36, (2008).

15. Rutter, M., Moffitt, T. E. & Caspi, A. Gene-environment interplay and psychopathology: Multiple varieties but real effects. Journal of Child Psychology and Psychiatry and Allied Disciplines vol. 47 Preprint at 10.1111/j.1469-7610.2005.01557.x (2006).

16. Claessens, S. E. F. et al. Development of individual differences in stress responsiveness: An overview of factors mediating the outcome of early life experiences. Psychopharmacology vol. 214 Preprint at 10.1007/s00213-010-2118-y (2011).

17. Balogh, S. A. & Wehner, J. M. Inbred mouse strain differences in the establishment of long-term fear memory. Behavioural Brain Research 140, (2003).

18. Bryant, C. D. et al. Behavioral differences among C57BL/6 substrains: Implications for transgenic and knockout studies. J Neurogenet 22, (2008).

19. Camp, M. C. et al. Genetic strain differences in learned fear inhibition associated with variation in neuroendocrine, autonomic, and amygdala dendritic phenotypes. Neuropsychopharmacology 37, (2012).

20. Cazares, V. A. et al. Environmental variables that ameliorate extinction learning deficits in the 129S1/SvlmJ mouse strain. Genes Brain Behav 18, (2019).

21. Eltokhi, A., Kurpiers, B. & Pitzer, C. Behavioral tests assessing neuropsychiatric phenotypes in adolescent mice reveal strain- and sex-specific effects. Sci Rep 10, (2020).

22. Hefner, K. et al. Impaired fear extinction learning and cortico-amygdala circuit abnormalities in a common genetic mouse strain. Journal of Neuroscience 28, (2008).

23. Owen, E. H., Logue, S. F., Rasmussen, D. L. & Wehner, J. M. Assessment of learning by the Morris water task and fear conditioning in inbred mouse strains and F1 hybrids: Implications of genetic background for single gene mutations and quantitative trait loci analyses. Neuroscience 80, (1997).

24. Park, K. & Chung, C. H. Differential Alterations in Cortico-Amygdala Circuitry in Mice with Impaired Fear Extinction. Mol Neurobiol 57, (2020).

25. Radulovic, J., Kammermeier, J. & Spiess, J. Generalization of fear responses in C57BL/6N mice subjected to one- trial foreground contextual fear conditioning. Behavioural Brain Research 95, (1998).

26. Ryherd, G. L., Bunce, A. L., Edwards, H. A., Baumgartner, N. E. & Lucas, E. K. Sex differences in avoidance behavior and cued threat memory dynamics in mice: Interactions between estrous cycle and genetic background. Horm Behav 156, (2023).

27. Jarome, T. J., Kwapis, J. L., Nye, S. H. & Helmstetter, F. J. Introgression of brown Norway chromosome 1 onto the fawn hooded hypertensive background rescues long-term fear memory deficits. Behav Genet 40, (2010).

28. Stöhr, T. et al. Lewis/Fischer rat strain differences in endocrine and behavioural responses to environmental challenge. Pharmacol Biochem Behav 67, (2000).

29. Jung, S. H. et al. Strain Differences in Responsiveness to Repeated Restraint Stress Affect Remote Contextual Fear Memory and Blood Transcriptomics. Neuroscience 444, (2020).

30. López-Aumatell, R. et al. A27 DIFFERENCES IN CLASSICAL FEAR CONDITIONING AND FEAR-POTENTIATED STARTLE BETWEEN THE ROMAN RAT STRAINS. Behavioural Pharmacology 16, (2005).

31. Schaap, M. W. H. et al. Nociception and conditioned fear in rats: Strains matter. PLoS One 8, (2013).

32. Cook, M. N., Bolivar, V. J., McFadyen, M. P. & Flaherty, L. Behavioral differences among 129 substrains: Implications for knockout and transgenic mice. Behavioral Neuroscience 116, (2002).

33. Wotjak, C. T. C57BLack/BOX? The importance of exact mouse strain nomenclature. Trends in Genetics vol. 19 Preprint at 10.1016/S0168-9525(02)00049-5 (2003).

34. Kiselycznyk, C. & Holmes, A. All (C57BL/6) mice are not created equal. Frontiers in Neuroscience Preprint at 10.3389/fnins.2011.00010 (2011).

35. Bothe, G. W. M., Bolivar, V. J., Vedder, M. J. & Geistfeld, J. G. Genetic and behavioral differences among five inbred mouse strains commonly used in the production of transgenic and knockout mice. Genes Brain Behav 3, (2004).

36. Siegmund, A., Langnaese, K. & Wotjak, C. T. Differences in extinction of conditioned fear in C57BL/6 substrains are unrelated to expression of α-synuclein. Behavioural Brain Research 157, (2005).

37. Siegmund, A. & Wotjak, C. T. A mouse model of posttraumatic stress disorder that distinguishes between conditioned and sensitised fear. J Psychiatr Res 41, (2007).

38. Stiedl, O. et al. Strain and substrain differences in context- and tone-dependent fear conditioning of inbred mice. Behavioural Brain Research 104, (1999).

39. Callaghan, B. L. & Richardson, R. The effect of adverse rearing environments on persistent memories in young rats: Removing the brakes on infant fear memories. Transl Psychiatry 10.1038/tp.2012.65 (2012) doi: 10.1038/tp.2012.65.

40. Gao, Y., Raine, A., Venables, P. H., Dawson, M. E. & Mednick, S. A. The development of skin conductance fear conditioning in children from ages 3 to 8 years. Dev Sci 13, (2010).

41. Gogolla, N., Caroni, P., Lüthi, A. & Herry, C. Perineuronal nets protect fear memories from erasure. Science (1979) 10.1126/science.1174146 (2009) doi: 10.1126/science.1174146.

42. Kim, J. H. & Richardson, R. A developmental dissociation in reinstatement of an extinguished fear response in rats. Neurobiol Learn Mem 10.1016/j.nlm.2007.03.004 (2007) doi: 10.1016/j.nlm.2007.03.004.

43. Kim, J. H. & Richardson, R. A developmental dissociation of context and GABA effects on extinguished fear in rats. Behavioral Neuroscience 121, (2007).

44. Park, C. H. J., Ganella, D. E. & Kim, J. H. Juvenile female rats, but not male rats, show renewal, reinstatement, and spontaneous recovery following extinction of conditioned fear. Learning and Memory 24, 630–636 (2017).

45. Josselyn, S. A. & Frankland, P. W. Infantile amnesia: A neurogenic hypothesis. Learning and Memory vol. 19 Preprint at 10.1101/lm.021311.110 (2012).

46. Alberini, C. M. & Travaglia, A. Infantile amnesia: A critical period of learning to learn and remember. Journal of Neuroscience 37, 5783–5795 (2017).

47. Ramsaran, A. I., Schlichting, M. L. & Frankland, P. W. The ontogeny of memory persistence and specificity. Dev Cogn Neurosci 36, (2018).

48. Callaghan, B. L., Li, S. & Richardson, R. The elusive engram: What can infantile amnesia tell us about memory? Trends Neurosci 37, 47–53 (2014).

49. Samifanni, R. et al. Developmental emergence of persistent memory for contextual and auditory fear in mice. Learning and Memory 10.1101/LM.053471.121 (2021) doi: 10.1101/LM.053471.121.

50. Yap, C. S. L. & Richardson, R. Extinction in the developing rat: An examination of renewal effects. Dev Psychobiol 49, (2007).

51. Park, C. H. J., Ganella, D. E. & Kim, J. H. Context fear learning and renewal of extinguished fear are dissociated in juvenile female rats. Dev Psychobiol 62, (2020).

52. Park, C. H. J., Ganella, D. E. & Kim, J. H. Juvenile female rats, but not male rats, show renewal, reinstatement, and spontaneous recovery following extinction of conditioned fear. Learning and Memory 24, 630–636 (2017).

53. Kim, J. H. & Richardson, R. A developmental dissociation in reinstatement of an extinguished fear response in rats. Neurobiol Learn Mem 10.1016/j.nlm.2007.03.004 (2007) doi: 10.1016/j.nlm.2007.03.004.

54. Kim, J. H. & Richardson, R. A developmental dissociation of context and GABA effects on extinguished fear in rats. Behavioral Neuroscience 121, (2007).

55. Gogolla, N., Caroni, P., Lüthi, A. & Herry, C. Perineuronal nets protect fear memories from erasure. Science (1979) 10.1126/science.1174146 (2009) doi: 10.1126/science.1174146.

56. Botterill, J. J. et al. Dorsal peduncular cortex activity modulates affective behavior and fear extinction in mice. Neuropsychopharmacology 49, (2024).

57. DeNardo, L. A. et al. Temporal evolution of cortical ensembles promoting remote memory retrieval. Nat Neurosci 22, (2019).

58. Matos, M. R. et al. Memory strength gates the involvement of a CREB-dependent cortical fear engram in remote memory. Nat Commun 10, (2019).

59. Lee, J. H., Kim, W. Bin, Park, E. H. & Cho, J. H. Neocortical synaptic engrams for remote contextual memories. Nat Neurosci 26, (2023).

60. Kitamura, T. et al. Engrams and circuits crucial for systems consolidation of a memory. Science (1979) 356, (2017).

61. Samifanni, R. et al. Developmental emergence of persistent memory for contextual and auditory fear in mice. Learning and Memory 28, 414–421 (2021).

62. Golbabaei, A., Josselyn, S. & Frankland, P. PV-dependent reorganization of prelimbic cortex sub-engrams during systems consolidation. Neuron (2025).

63. Ganella, D. E., Barendse, M. E. A., Kim, J. H. & Whittle, S. Prefrontal-amygdala connectivity and state anxiety during fear extinction recall in adolescents. Front Hum Neurosci 11, (2017).

64. Zeiss, C. J. Comparative Milestones in Rodent and Human Postnatal Central Nervous System Development. Toxicol Pathol 49, (2021).

65. Arruda-Carvalho, M., Wu, W. C., Cummings, K. A. & Clem, R. L. Optogenetic examination of prefrontal-amygdala synaptic development. Journal of Neuroscience 37, 2976–2985 (2017).

66. Baker, K. D., Gray, A. R. & Richardson, R. The development of perineuronal nets around parvalbumin GABAergic neurons in the medial prefrontal cortex and basolateral amygdala of rats. Behavioral Neuroscience 10.1037/bne0000203 (2017) doi: 10.1037/bne0000203.

67. Chareyron, L. J., Lavenex, P. B. & Lavenex, P. Postnatal development of the amygdala: A stereological study in rats. Journal of Comparative Neurology 10.1002/cne.23132 (2012) doi: 10.1002/cne.23132.

68. Glantz, L. A., Gilmore, J. H., Hamer, R. M., Lieberman, J. A. & Jarskog, L. F. Synaptophysin and postsynaptic density protein 95 in the human prefrontal cortex from mid-gestation into early adulthood. Neuroscience 10.1016/j.neuroscience.2007.06.036 (2007) doi: 10.1016/j.neuroscience.2007.06.036.

69. Koss, W. A., Belden, C. E., Hristov, A. D. & Juraska, J. M. Dendritic remodeling in the adolescent medial prefrontal cortex and the basolateral amygdala of male and female rats. Synapse 10.1002/syn.21716 (2014) doi: 10.1002/syn.21716.

70. Kroon, T., van Hugte, E., van Linge, L., Mansvelder, H. D. & Meredith, R. M. Early postnatal development of pyramidal neurons across layers of the mouse medial prefrontal cortex. Sci Rep 9, (2019).

71. Ryan, S. J., Ehrlich, D. E. & Rainnie, D. G. Morphology and dendritic maturation of developing principal neurons in the rat basolateral amygdala. Brain Struct Funct 221, (2016).

72. Tottenham, N. & Gabard-Durnam, L. J. The developing amygdala: a student of the world and a teacher of the cortex. Current Opinion in Psychology vol. 17 Preprint at 10.1016/j.copsyc.2017.06.012 (2017).

73. Zhang, Z. W. Maturation of Layer V Pyramidal Neurons in the Rat Prefrontal Cortex: Intrinsic Properties and Synaptic Function. J Neurophysiol 91, (2004).

74. Zimmermann, K. S., Richardson, R. & Baker, K. D. Maturational changes in prefrontal and amygdala circuits in adolescence: implications for understanding fear inhibition during a vulnerable period of development. Brain Sci 9, 65 (2019).

75. Travaglia, A., Bisaz, R., Cruz, E. & Alberini, C. M. Developmental changes in plasticity, synaptic, glia and connectivity protein levels in rat dorsal hippocampus. Neurobiol Learn Mem 135, (2016).

76. Li, L., Gao, X. & Zhou, Q. Absence of fear renewal and functional connections between prefrontal cortex and hippocampus in infant mice. Neurobiol Learn Mem 152, (2018).

77. Ramsaran, A. I. et al. A sensitive period for the development of episodic-like memory in mice. Current Biology 35, 2032–2048.e3 (2025).

78. Ramsaran, A. I. et al. A shift in the mechanisms controlling hippocampal engram formation during brain maturation. Science (1979) 380, (2023).

79. Cho, J. H., Deisseroth, K. & Bolshakov, V. Y. Synaptic encoding of fear extinction in mPFC-amygdala circuits. Neuron 80, (2013).

80. Feng, P., Zheng, Y. & Feng, T. Resting-state functional connectivity between amygdala and the ventromedial prefrontal cortex following fear reminder predicts fear extinction. Soc Cogn Affect Neurosci 11, (2016).

81. Kim, J. H. & Richardson, R. The effect of temporary amygdala inactivation on extinction and reextinction of fear in the developing rat: Unlearning as a potential mechanism for extinction early in development. Journal of Neuroscience 10.1523/JNEUROSCI.4736-07.2008 (2008) doi: 10.1523/JNEUROSCI.4736-07.2008.

82. Marek, R., Sun, Y. & Sah, P. Neural circuits for a top-down control of fear and extinction. Psychopharmacology vol. 236 Preprint at 10.1007/s00213-018-5033-2 (2019).

83. Vouimba, R. M. & Maroun, M. Learning-induced changes in mpfc-bla connections after fear conditioning, extinction, and reinstatement of fear. Neuropsychopharmacology 36, (2011).

84. Marek, R. et al. Hippocampus-driven feed-forward inhibition of the prefrontal cortex mediates relapse of extinguished fear. Nat Neurosci 21, (2018).

85. Plas, S. L. et al. Neural circuits for the adaptive regulation of fear and extinction memory. Frontiers in Behavioral Neuroscience vol. 18 Preprint at 10.3389/fnbeh.2024.1352797 (2024).

86. Bechard, A. & Mason, G. Leaving home: A study of laboratory mouse pup independence. Appl Anim Behav Sci 125, (2010).

87. Gee, D. G. et al. A developmental shift from positive to negative connectivity in human amygdala-prefrontal circuitry. Journal of Neuroscience 33, (2013).

88. Gee, D. G. et al. Early developmental emergence of human amygdala-prefrontal connectivity after maternal deprivation. Proc Natl Acad Sci U S A 10.1073/pnas.1307893110 (2013) doi: 10.1073/pnas.1307893110.

89. Wiggins, J. L. et al. Age-related effect of serotonin transporter genotype on amygdala and prefrontal cortex function in adolescence. Hum Brain Mapp 35, (2014).

90. Johnson, F. K. et al. Amygdala hyper-connectivity in a mouse model of unpredictable early life stress. Translational Psychiatry vol. 8 Preprint at 10.1038/s41398-018-0092-z (2018).

91. Honeycutt, J. A. et al. Altered corticolimbic connectivity reveals sex-specific adolescent outcomes in a rat model of early life adversity. Elife 10.7554/eLife.52651 (2020) doi: 10.7554/eLife.52651.

92. Baker, K. D. & Richardson, R. Forming competing fear learning and extinction memories in adolescence makes fear difficult to inhibit. Learning and Memory 22, (2015).

93. Baker-Andresen, D., Flavell, C. R., Li, X. & Bredy, T. W. Activation of BDNF signaling prevents the return of fear in female mice. Learning and Memory 10.1101/lm.029520.112 (2013) doi: 10.1101/lm.029520.112.

94. Ganella, D. E. et al. Aripiprazole facilitates extinction of conditioned fear in adolescent rats. Front Behav Neurosci 11, (2017).

95. McCallum, J., Kim, J. H. & Richardson, R. Impaired extinction retention in adolescent rats: Effects of d-cycloserine. Neuropsychopharmacology 35, (2010).

96. Pattwell, S. S. et al. Altered fear learning across development in both mouse and human. Proc Natl Acad Sci U S A 10.1073/pnas.1206834109 (2012) doi: 10.1073/pnas.1206834109.

97. Pattwell, S. S. et al. Dynamic changes in neural circuitry during adolescence are associated with persistent attenuation of fear memories. Nat Commun 7, (2016).

98. Premachandran, H., Wilkin, J. & Arruda-Carvalho, M. Minimizing variability in developmental fear studies in mice: Toward Improved Replicability in the Field. Curr Protoc 4, e1040 (2024).

99. Kim, J. H., Li, S. & Richardson, R. Immunohistochemical analyses of long-term extinction of conditioned fear in adolescent rats. Cerebral Cortex 21, (2011).

100. Gerhard, D. M. & Meyer, H. C. Extinction trial spacing across days differentially impacts fear regulation in adult and adolescent male mice. Neurobiol Learn Mem 186, (2021).

101. McCallum, J., Kim, J. H. & Richardson, R. Impaired extinction retention in adolescent rats: Effects of d-cycloserine. Neuropsychopharmacology 35, (2010).

102. Gerhard, D. M., Tse, N., Lee, F. S. & Meyer, H. C. Developmental age and fatty acid amide hydrolase genetic variation converge to mediate fear regulation in female mice. Dev Psychobiol 65, (2023).

103. An, X. L. et al. Strain and sex Differences in Anxiety-Like and Social Behaviors in C57Bl/6J and BALB/cJ Mice. Exp Anim 60, (2011).

104. Ryherd, G. L., Bunce, A. L., Edwards, H. A., Baumgartner, N. E. & Lucas, E. K. Sex differences in avoidance behavior and cued threat memory dynamics in mice: Interactions between estrous cycle and genetic background. Horm Behav 156, (2023).

105. Võikar, V., Kõks, S., Vasar, E. & Rauvala, H. Strain and gender differences in the behavior of mouse lines commonly used in transgenic studies. Physiol Behav 72, (2001).

106. McCutcheon, T. B., Simenson-Braun, S. M. & Richardson, R. Accelerated maturation of fear regulation systems in infant rats following early life inflammation. Neurobiol Learn Mem 222, 108109 (2025).

107. Bell, M. R. Comparing postnatal development of gonadal hormones and associated social behaviors in rats, mice, and humans. Endocrinology 159, (2018).

108. Keeley, R. J., Trow, J. & McDonald, R. J. Strain and sex differences in puberty onset and the effects of THC administration on weight gain and brain volumes. Neuroscience 305, (2015).

109. Nelson, J. F., Karelus, K., Felicio, L. S. & Johnson, T. E. Genetic influences on the timing of puberty in mice. Biol Reprod 42, (1990).

110. Piekarski, D. J., Boivin, J. R. & Wilbrecht, L. Ovarian Hormones Organize the Maturation of Inhibitory Neurotransmission in the Frontal Cortex at Puberty Onset in Female Mice. Current Biology 10.1016/j.cub.2017.05.027 (2017) doi: 10.1016/j.cub.2017.05.027.

111. Premachandran, H., Zhao, M. & Arruda-Carvalho, M. Sex Differences in the Development of the Rodent Corticolimbic System. Frontiers in Neuroscience Preprint at 10.3389/fnins.2020.583477 (2020).

112. Chang, Y. J. et al. Estrogen modulates sexually dimorphic contextual fear extinction in rats through estrogen receptor β. Hippocampus 19, (2009).

113. Colón, L., Peru, E., Zuloaga, D. G. & Poulos, A. M. Contributions of gonadal hormones in the sex-specific organization of context fear learning. PLoS One 18, (2023).

114. Cover, K. K., Maeng, L. Y., Lebrón-Milad, K. & Milad, M. R. Mechanisms of estradiol in fear circuitry: Implications for sex differences in psychopathology. Translational Psychiatry Preprint at 10.1038/tp.2014.67 (2014).

115. Crestani, A. P. et al. Adolescent Female Rats Undergo Full Systems Consolidation of an Aversive Memory, While Males of the Same Age Fail to Discriminate Contexts. Behavioral Neuroscience 136, (2022).

116. Gupta, R. R., Sen, S., Diepenhorst, L. L., Rudick, C. N. & Maren, S. Estrogen modulates sexually dimorphic contextual fear conditioning and hippocampal long-term potentiation (LTP) in rats. Brain Res 888, (2001).

117. Jasnow, A. M., Schulkin, J. & Pfaff, D. W. Estrogen facilitates fear conditioning and increases corticotropin-releasing hormone mRNA expression in the central amygdala in female mice. Horm Behav 49, (2006).

118. Lebron-Milad, K. & Milad, M. R. Sex differences, gonadal hormones and the fear extinction network: implications for anxiety disorders. Biol Mood Anxiety Disord 2, (2012).

119. McDermott, C. M., Liu, D. & Schrader, L. A. Role of gonadal hormones in anxiety and fear memory formation and inhibition in male mice. Physiol Behav 105, (2012).

120. Milad, M. R., Igoe, S. A., Lebron-Milad, K. & Novales, J. E. Estrous cycle phase and gonadal hormones influence conditioned fear extinction. Neuroscience 164, 887–895 (2009).

121. Perry, C. et al. Assessment of conditioned fear extinction in male and female adolescent rats. Psychoneuroendocrinology 116, 104670 (2020).

122. Bryant, C. D. et al. Behavioral differences among C57BL/6 substrains: Implications for transgenic and knockout studies. J Neurogenet 22, (2008).

